# The Role of Phylogenetic Structure on Canidae Lineage Distribution and Trait Variation across the Globe

**DOI:** 10.1101/2023.07.08.548220

**Authors:** Geographical patterns of Canidae phylogenetic information, Lucas Marafina Vieira Porto, Arielli Fabricio Machado

## Abstract

**Aim:** Understanding the spatial structuring of ecological communities requires us to consider the interplay between evolutionary history and environmental factors. In this study, we investigated the influence of Canidae phylogenetic structure on the distribution and trait patterns of lineages across the planet.

**Location:** Americas, Africa, Eurasia.

**Time period:** 12 million years ago – present.

**Major taxa studied:** Canidae.

**Methods:** Using distribution data and phylogenetic information for 37 Canidae species, we employed phylogenetic fuzzy-weighting to compare assemblages based on their phylogenetic similarity.

**Results:** Our results revealed distinct global patterns of body size, body mass, range size, habitat type, and evolutionary distinctiveness among lineages. We also identified the shared contributions of phylogenetic structure and temperature to trait variation using variance partitioning analysis. The PCPS axes highlighted the influence of phylogenetic relationships on Canidae assemblages, particularly in South America.

**Main conclusions:** Notably, the study challenges the applicability of Bergmann’s and Rapoport’s rules in explaining canids’ sizes and range sizes geographic variation across continents, emphasizing the importance of phylogenetic information. The unique diversification history of Canidae in South America and Africa and their diverse environmental conditions likely contribute to the observed trait patterns that make both continents so distinguished when compared to N. America and Eurasia. Our findings underscore the need to incorporate phylogenetic information in models assessing trait variation across geographic scales for unbiased estimations.

## INTRODUCTION

Understanding how evolutionary history and environmental factors drive the spatial structuring of ecological communities is a fundamental and exciting goal in ecology. The distribution of species is shaped by a complex interplay between biotic and abiotic factors over time (Webb et al. 2002; Westoby 2006). For a long time, studies on the variation of species traits across geography have traditionally focused on understanding the relationship between those traits and the environmental gradient across space (Rodríguez et al. 2008; Šizling et al. 2009). However, the phylogenetic information is a crucial aspect to assess the variation in the life-history strategies across the planet, as one cannot consider species as independent units (Wiens and Donoghue 2004). Macroevolutionary approaches developed over the last decades took this phylogenetic issue into account, making it possible to explore the association among traits and spatial gradients over time and across lineages (Wiens et al. 2010; Pennell et al. 2014). Thus, to have a more realistic assessment of the mechanisms underneath the distribution of both species and their traits, the evolutionary history traced by lineages across space must be considered.

Patterns of biogeography have been extensively explored in vertebrates, revealing intriguing insights into the distribution of species across geographic regions. Two prominent biogeographic patterns in vertebrates are known as Bergmann’s rule and Rapoport’s rule. Bergmann’s rule states that, among closely related lineages, species tend to be larger in colder climates and smaller in warmer climates (Bergmann 1847). This pattern has been observed in various groups of vertebrates, including mammals (Clauss et al. 2013), birds (Jirinec et al. 2021), and reptiles (Lakin et al. 2020). On the other hand, Rapoport’s rule proposes that species’ latitudinal ranges tend to be broader at higher latitudes compared to lower latitudes, which suggests that species occupying higher latitudes experience greater environmental variability, leading to a wider distribution to encompass a range of ecological conditions (Stevens 1989). Although both rules have been criticized in the past as other factors can also influence such patterns, they hold well in mammals and birds and can be considered valid ecological generalizations in these endothermic groups (Meiri and Dayan 2003; Salewski and Watt 2017).

Furthermore, the growing accessibility of phylogenetic data has sparked a surge of interest in studying how phylogenetic information is distributed across space (i.e., Phylogeography). Patterns of Phylogeography are helpful to understand if co-occurring species into assemblages are more phylogenetically closed or distant than expected by chance, and how the traits they share can shed light on the mechanisms that assembled such area (Webb et al. 2002; Ackerly et al. 2006). Understanding how species diversity is structured in space and time remains one of the main challenges in the fields of ecology and evolution, and is considered one of the 100 fundamental ecological questions (Sutherland et al. 2013).

The Canidae family presents an ideal opportunity for such investigations on how traits and phylogenetic information are distributed in space. Canids are found on all continents, except for Antarctica, making them geographically diverse (Wang and Tedford 2008; Wilson and Mittermeier 2009). As canids inhabit several environmental settings, they exhibit a wide range of attributes, reflecting the evolutionary history of species (Wilson and Mittermeier 2009). Additionally, the phylogenetic tree of canids is highly resolved, providing a solid foundation for analyzing the evolutionary relationships within the clade (Porto et al. 2019).

In this study, we explored the influence of Canidae phylogenetic structure on how lineages are distributed across the planet. We also assessed patterns of behavioral and morphological traits over space. To test such patterns (i.e., Bergmann’s and Rapoport’s rules), commonly used metrics such as the Net Relatedness Index (NRI) (Webb 2000; Webb et al. 2002), which indicates the degree to which species within the community are more or less closely related than expected by chance, can impose a problem, as communities that contain totally distinct phylogenetic clades can rise similar levels of phylogenetic clustering or overdispersion. Thus, we used phylogenetic fuzzy-weighting, which allows us to define local assemblages by their phylogeny-weighted species composition (Duarte 2011). Such an approach makes it possible to compare assemblages in terms of their phylogenetic similarity.

## MATERIAL AND METHODS

### Sampling data and phylogenetic tree

We used the phylogeny from Porto et al. (2019) that presents 37 Canidae species (Figure S1) (all the 36 extant canids on earth plus one recently extinct, *Dusicyon australis*, in 1867) (IUCN 2020). We used this phylogeny because it presents robust relationships among lineages, being constructed under a Bayesian framework using 22 mitochondrial and molecular markers, besides more than 60 osteological traits. The phylogeny of Canidae presents three distinct clades: True-wolves, Cerdocyonina, and Vulpini (Wang and Tedford 2008; Tedford et al. 2009), which we will call hereafter as wolves, South American canids, and foxes, respectively (Figure S1).

Distribution data for 37 Canidae species were obtained from IUCN distribution polygons (IUCN 2020) (Figure S2). We used the IUCN distribution data since these maps are created and verified by specialists based on occurrence records and restrict species occurrences to areas with the presumably suitable habitat where the species is known, following a precautionary principle to guide conservation efforts described in the Mapping Standards and Data Quality for IUCN Red List Spatial Data (IUCN 2021). Although these maps were designed for conservation purposes, they have been shown to be an important source of information in macroecological studies (Belmaker and Jetz 2015; Vilela and Villalobos 2015; Maestri et al. 2016; Gonçalves-Souza et al. 2021).

To explore how biotic and abiotic patterns are associated with the geographic distribution of the phylogenetic information of canids, we mapped how traits like body size (cm), body weight (kg), range size (km^2^), and evolutionary distinctiveness among lineages (Myr) are distributed across the globe. These traits were obtained from *ADW* and *PanTHERIA* (Jones et al. 2009; Myers et al. 2018). In addition, mean global temperature (°C) and percentage of vegetation cover (0 – 100%) were also used here (IUCN 2020). The evolutionary distinctiveness was measured using the *picante* package (Kembel et al. 2010) in R (R Development Core Team 2023). To observe how traits are distributed across assemblages, we used them to compute the Community Weighted Means (CWMs) using the package *FD* (Laliberté et al. 2022).

### Data analysis

We Computed the phylogenetic distance between tips using the method patristic by the *distTips* function of the R package *adephylo* 1.1-13 (Jombart et al. 2010).

Distribution polygons for each species were converted into raster layers containing grids of cells of 1 x 1° (∼ 110 x 110 km) and extract the coordinates of occurrence for each species using the R package *maptools* 1.1-7 (Bivand and Lewin-Koh 2023) and *raster* 3.6-20 (Hijmans and Etten 2012), creating a matrix of presence and absence for all species (i.e. a community data, with species as columns and sampling units as rows).

We generated Principal Coordinates of Phylogenetic Structure (PCPS) (Duarte 2011) using the *pcps* function of the R Package *PCPS* 1.0.7 (Debastiani 2020) and *organize*.*syncsa* of the R Package *SYNCSA* 1.3.3 (Debastiani and Pillar 2012). The phylogenetic information (PCPS) and all other traits used here were projected onto a species-by-plot matrix into geographic space using the *matrix*.*t* function of the R package *SYNCSA* using the community matrix. This procedure generated a matrix describing each plot (grid-cell) by each trait, environmental variable, or phylogenetic information. The PCPS with higher eigenvalues usually describes phylogenetic gradients more associated with the deepest nodes of the phylogenetic tree, while the PCPS with smaller eigenvalues describes finer phylogenetic gradients (Duarte et al. 2012).

The phylogenetic signal in body size, body weight, habitat type, and range size across all 37 canids was tested using the PSR curve (Diniz Filho et al. 2012). PSR curve tests the strength of the phylogenetic signal in a trait allowing us to compare the observed signal with expectations from either the Brownian or the OU model. If a trait has evolved under a Brownian motion model, a linear relationship (45° line) is expected. Deviations from this line indicate either a faster (higher R^2^) or slower (lower R^2^) evolution of the trait than that expected under Brownian motion (Diniz Filho et al. 2012). All the analyses were performed using the R software (R Development Core Team 2023).

We applied a variance partitioning analysis (VPA) (Borcard et al. 1992) to estimate the relative contribution of phylogenetic structure (PCPS) and environment (temperature) to all Canidae traits used here as response variables (body size, body mass, habitat type, range size, and evolutionary distinctiveness). We only used two first axes of PCPS during this step. VPA was conducted in R software using the *vegan* package (Oksanen et al. 2013).

## RESULTS

The phylogenetic signal for body size, body weight, habitat type, and range size produced PSR areas of -0.03, -0.15, -0.18, and -0.25, respectively. All traits presented *P* values < 0.05 (Figure 1). The PSR areas of all traits are nearest to a Brownian motion model of evolution.

**Figure 1.**
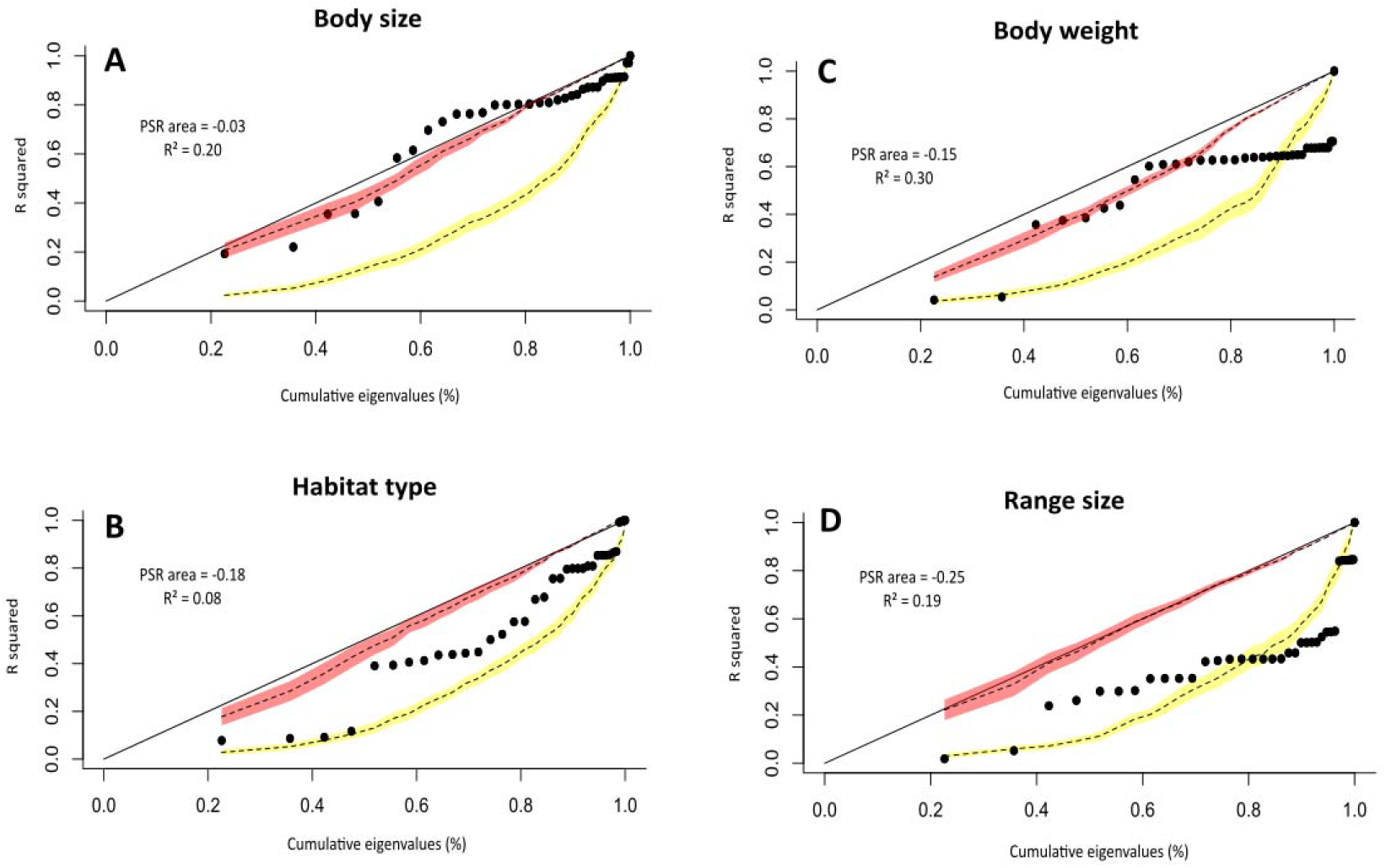
Phylogenetic signal-representation (PSR) curve on body size, body weight, habitat type, and range size. Black dots show the PSR curve for the values of each trait. The dashed line with a red shadow shows the expected curve under Brownian motion. The dashed line with a yellow shadow indicates the null expectation.

CWMs plot across the globe (Figure 2) reveals distinct patterns. Canid body mass (Figure 2A) shows a higher concentration in the Northern Hemisphere, while larger body sizes are observed at higher latitudes and on islands such as New Zealand, western Greenland, and the Falkland Islands (Figure 2B). Regarding range size, species inhabiting the Southern Hemisphere exhibit smaller ranges compared to those in the Northern Hemisphere (Figure 2C). In addition, when we compare the range sizes of species that inhabit each hemisphere, it seems to be a lot of variation among the ones within the North Hemisphere, but the canids in the South of the globe present almost no variation in this trait.

**Figure 2.**
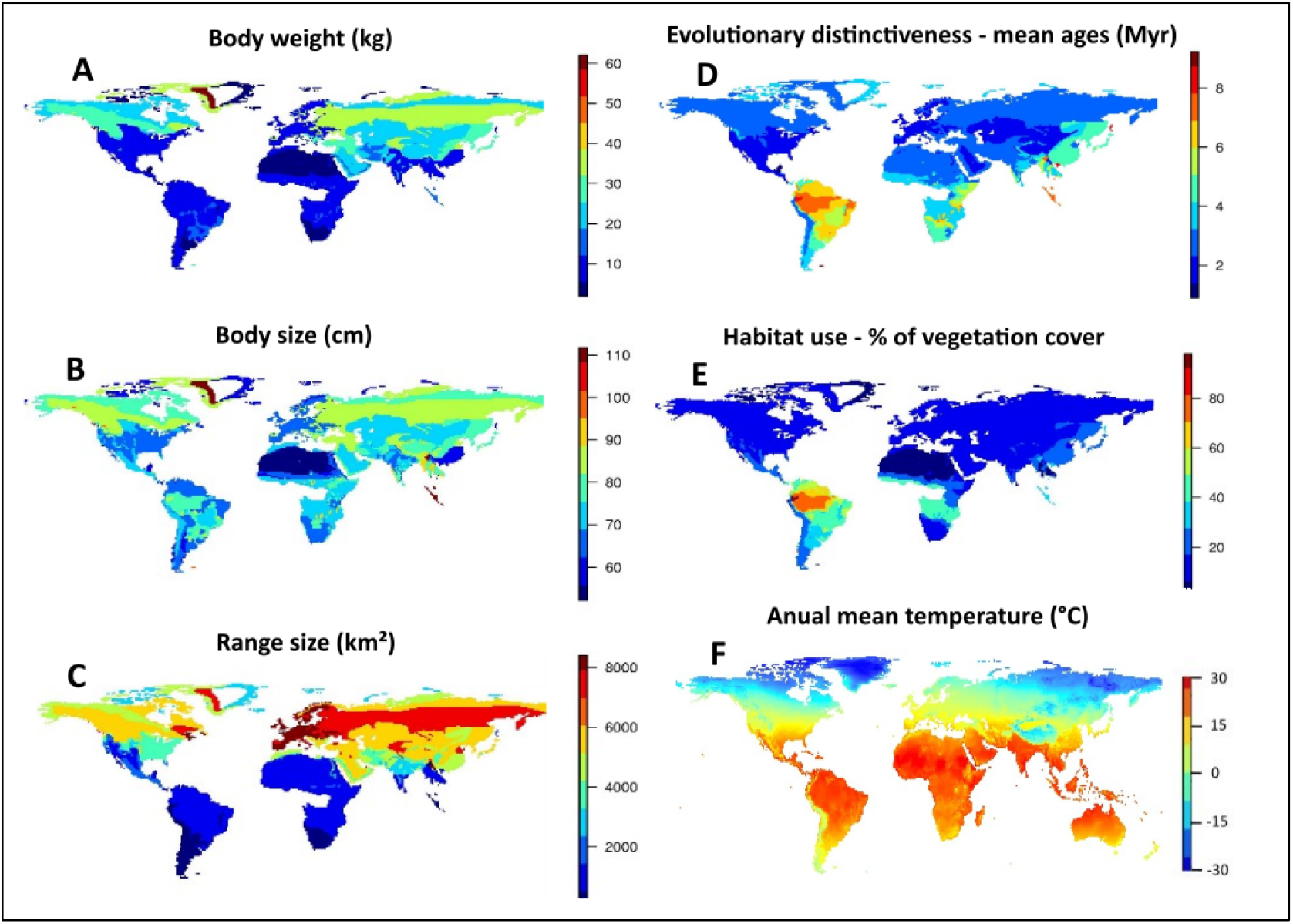
Geographic distribution of traits and environmental data used here based on CWM values.

Canidae lineages distributed near the Equator line are (on average) the oldest ones on the planet, presenting ages around 7 Myr, which can be observed across the tropical region of South America, Africa, and Asia (Figure 2D). Besides, South America presents a strong variation of ages compared to the rest of the planet. We can also notice that, within South America, lineages inhabiting the west of the Andes are younger compared to the ones on the East of the Andean Mountain. In terms of habitat use, canids with a stronger affinity for forests are predominantly found in South America, whereas other continents harbor species associated with open areas (Figure 2E). Furthermore, the association between the latitudinal gradient with body mass and body size of canids was weak (Figures S3A and S3B).

PCPS 1 and 2 are responsible for 36% and 21%, respectively, of the total variation of the phylogenetic information. Both PCPS axes revealed how the Canidae tribes (South American canids, wolves, and foxes) structure assemblages across the globe (Figure 3A). Values along PCPS 1 are responsible for mainly separating the endemic South American lineages from the tribes that inhabit the rest of the planet (wolves and foxes). The segregation of both cosmopolitan tribes is largely due to PCPS 2.

**Figure 3.**
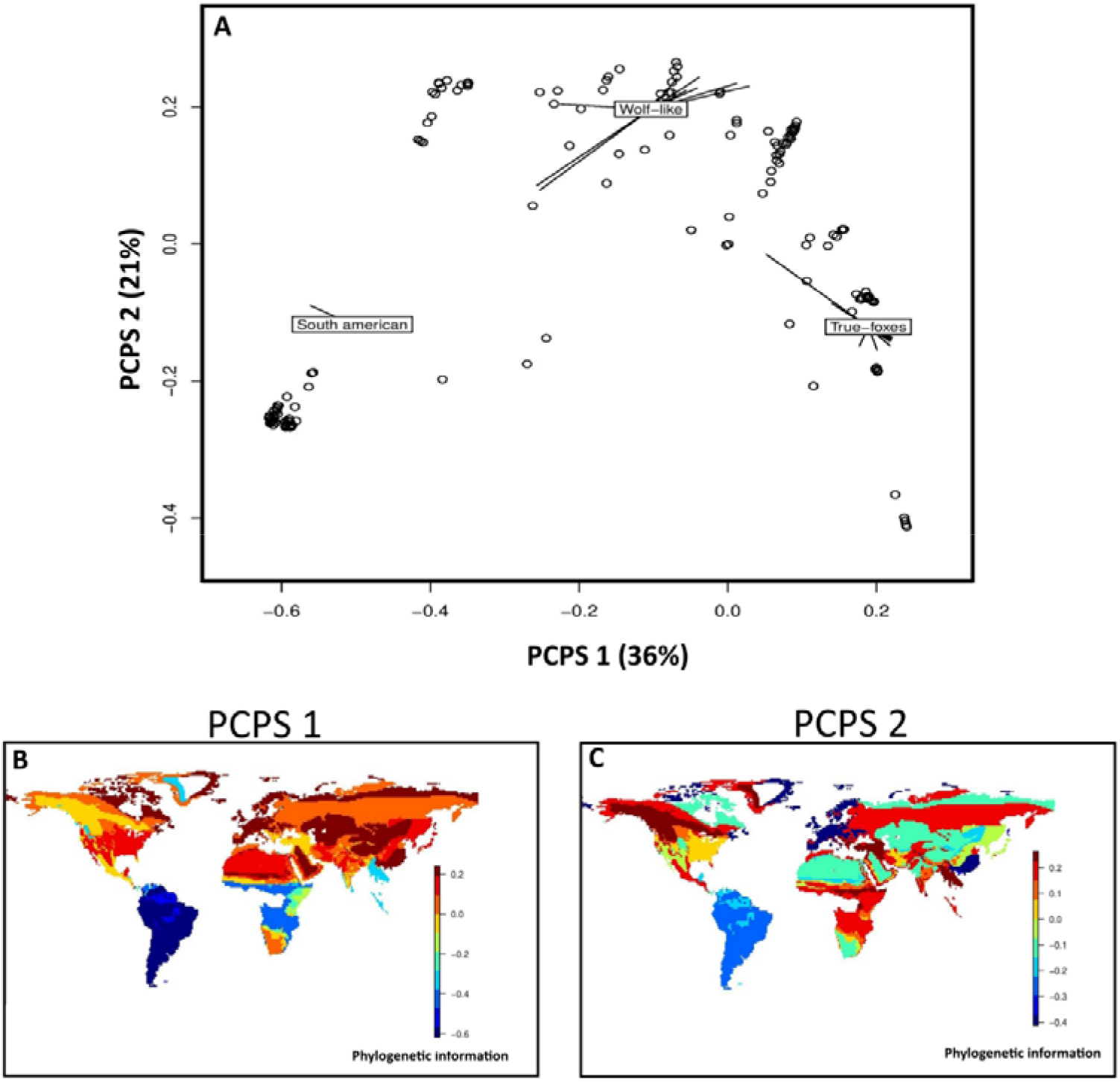
Information regarding the two main axes of the phylogenetic composition of Canidae. (A) Correlation plot among PCPS 1 and 2. The three tribes of Canidae are indicated, and circles are the sites used here. (B and C) The geographical pattern of PCPS 1 and 2 across the sites.

The geographical variation in the phylogenetic composition of assemblages is observed by the distribution of the PCPS 1 and 2 on the map (Figure 3B and 3C). PCPS 1 contrasts South America with other continents. But the variation of phylogenetic information among wolves and foxes can only be observed on assemblages using the PCPS 2.

Variance partitioning analysis indicated that the shared contribution of PCPS axes and temperature is essential to understand the geographic distribution patterns of body mass, body size, range size, evolutionary distinctiveness, and habitat type (Table 1).

**Table 1.** Variation partitioning results by site among our response variables against environmental variables and phylogenetic composition of each site as the explanatory variables.

| Response variables | PCPS 1 + PCPS 2 | Temperature | Shared contribution |  |
| --- | --- | --- | --- | --- |
| | Adj. $R^2$ | Adj. $R^2$ | Adj. $R^2$ | $P$ |
| Range size | 0.40 | 0.43 | 0.57 | $P < 0.05$ |
| Body size | 0.44 | 0.20 | 0.67 | $P<0.05$ |
| Body mass | 0.37 | 0.40 | 0.62 | $P<0.05$ |
| Evol. distinctiveness | 0.45 | 0.34 | 0.55 | $P<0.05$ |
| Habitat | 0.67 | 0.19 | 0.67 | $P<0.05$ |

## DISCUSSION

Our findings indicated that the shared effect of historical processes related to the distribution of Canidae lineages across the planet and the temperature gradient has significantly influenced the geographical variation in all the traits assessed here. The PCPS axes plotted on the map highlighted the strong influence of phylogenetic relationships among Canidae assemblages in South America compared to the rest of the continents. Such results emphasize the importance of considering the phylogenetic information of species present in a specific location as a crucial historical factor in explaining the variation in its traits across geographical scales, as long as such traits present a certain degree of phylogenetic conservation, which was our case here. Thus, not incorporating the phylogenetic information of species’ locations on models to evaluate patterns of trait variation across space may lead to biased estimations.

We presented results that reflect a direct implication for the use of Bergmann’s rule in understanding how species’ sizes vary across space. Although studies already demonstrated a trend followed by mammals that consists of an increase of body mass in higher latitudes (Blackburn and Hawkins 2004; Rodríguez et al. 2008), this trend is not presented by Canidae (Gohli and Voje 2016). Although the mean trait value maps present bigger and heavier canids in higher latitudes in the Northern Hemisphere, this did not happen in the Southern Hemisphere (Figures 2A and 2B), and both correlations are very weak for the whole distribution of Canidae (Figures S3A and S3B). We argue that neither the latitude nor temperature are enough to explain how canids vary their sizes across the planet, but the phylogenetic information is essential to this issue, as the combined effect of PCPS axes and temperature on body size and body mass (two traits that can reflect species’ sizes) is strong (Table 1). One possible explanation for the absence of a clear Bergmann’s rule effect, and the major influence of the phylogenetic information on trait geographic variation in canids is the process in which canids colonized and diversified into South America and Africa. The findings from Porto et al. (2023), on the unique diversification history of these lineages, shed light on the weak effect of Bergmann’s rule on this group. Their study revealed three major centers of diversification for Canidae over the past 12 million years: South America, Africa, and North America. However, unlike S. America and Africa, N. America experienced several dispersal events of its daughter lineages to Eurasia and also received some lineages originating from there, which could have resulted in a loss of its distinct “identity” compared to the other two diversification centers. We hypothesize that the greater endemism maintained by Africa and S. America (Porto et al. 2023), along with their highly diverse environmental conditions compared to the Northern Hemisphere (Rahbek 1997), might have contributed to the observed geographic pattern of size variation across the continents observed here.

Rapoport’s rule is another well-studied biogeographic pattern that fails to explain the overall geographic distribution of Canidae range sizes across the planet. In the Northern Hemisphere, Canids follow Rapoport’s rule, with larger ranges associated with higher latitudes (Figure 2C). However, in the Southern Hemisphere, a contrasting pattern emerges, where Canids exhibit smaller ranges at lower latitudes. Interestingly, even though the smaller latitudes in the Southern Hemisphere encompass less area compared to the larger latitudes in the Northern Hemisphere, this consistent pattern persists throughout South America and Africa. The combined effect of temperature and phylogenetic information becomes crucial in understanding the underlying mechanisms shaping range size patterns in these regions, where traditional biogeographic rules do not apply as neatly. Once again, it is likely that the way canids diversified within both continents and explored the distinct niches available acted in a distinct way to shape their range sizes. Larger tropical zones with less seasonality over the last 12 million years could help to understand this strong difference between the Northern and Southern Hemispheres (Hawkins and Diniz-Filho 2006; Willig and Presley 2018).

When examining the evolutionary distinctiveness among canids in South America (Figure 2D), an intriguing pattern emerges. We observed that the lineages to the west of the Andes are considerably more recent than those inhabiting the east of the mountains. This suggests that canids initially invaded South America through the eastern region, and later circumvented the Andes along their southern portion to reach the west. These findings align with the results from Chavez et al. (2022) and Porto et al. (2023), who also suggested an initial colonization of South America from the east, followed by the arrival of canids in the western Andes later. Thus, from a new perspective, our study provides further support for such major dispersal events.

The first PCPS captured the phylogenetic gradient responsible for splitting South American canids from wolves and foxes (capturing a phylogenetic gradient related to more intermediary nodes). The second PCPS captured a basal phylogenetic gradient, splitting wolves and foxes. The observed phylogenetic gradient along the distribution of Canidae (Figures 3B and 3C) played a crucial role in explaining the variation of other canid attributes in our study. Unlike other methods that assess assembly structure, such as Rao’s H (Rao 1982) and Net Relatedness Index (Webb et al. 2002), PCPS allowed us to explore the specific clades responsible for driving phylobetadiversity among sites. In addition, PCPS shed light on the geographic gradient in body mass, body size, habitat type, range size, and evolutionary distinctiveness of canids, but it also revealed a particularly intriguing pattern that distinguishes South America from other continents. Phylogenetic maps (Figures 3B and 3C) highlight the significance of canid diversification in South America as a major event for the clade. A single dispersal event from North to S. America led to the rise of a diverse array of endemic lineages in this region, while other continents, including Africa, received and still harbor species that originated and are distributed across more than one continent over the past millions of years (Porto et al. 2023). The evolutionary distinctiveness map demonstrates that the lineages in S. America are older than those in other continents (Figure 2D). Thus, we hypothesize that the unexplored fauna of South America, in the absence of major competitors at the time of canid arrival (Wang and Tedford 2008), was crucial in shaping this pattern, resulting in more generalist species in terms of both habitat and diet. Such generalist behavior enables species to adapt to changing environmental conditions and utilize a wider range of resources compared to specialists, which likely contributed to their long persistence over evolutionary time, which was already suggested by Naeem and Li (1997) and Mouillot et al. (2013).

In conclusion, our findings indicate that the phylogenetic composition of canid assemblages plays a crucial role in shaping the five traits of Canidae assessed here across different geographical regions. The geographic variation of Canidae traits is strongly influenced by the distribution of specific canid lineages within assemblages. These results emphasize the importance of considering the explicit context of lineage distribution when examining the geography of trait variation. Therefore, we indicated here that environmental factors alone, or even the latitudinal gradient are insufficient to explain the complex processes underlying trait variation in canids across the planet. We suggest that future investigations into S. America and Africa should be done in order for us to have a better understanding of why canids in both continents do not follow Bergmann’s and Rapoport’s rules.

## SUPPLEMENTARY MATERIAL

**Figure S1.**
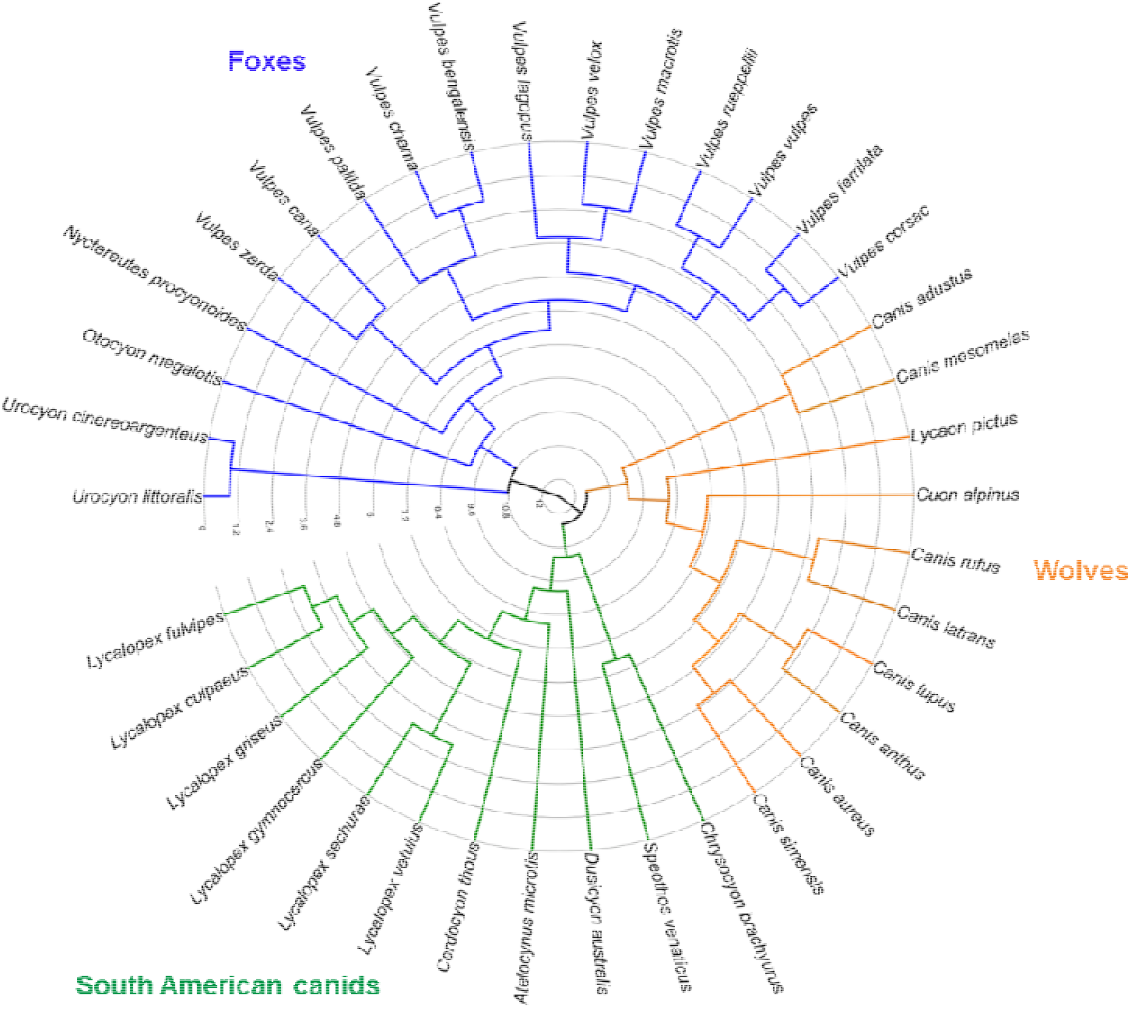
Phylogenetic tree from Porto et al. (2019) that was used during our study. The three major clades of S. American canids, wolves, and foxes are represented by green, orange and blue colors.

**Figure S2.**
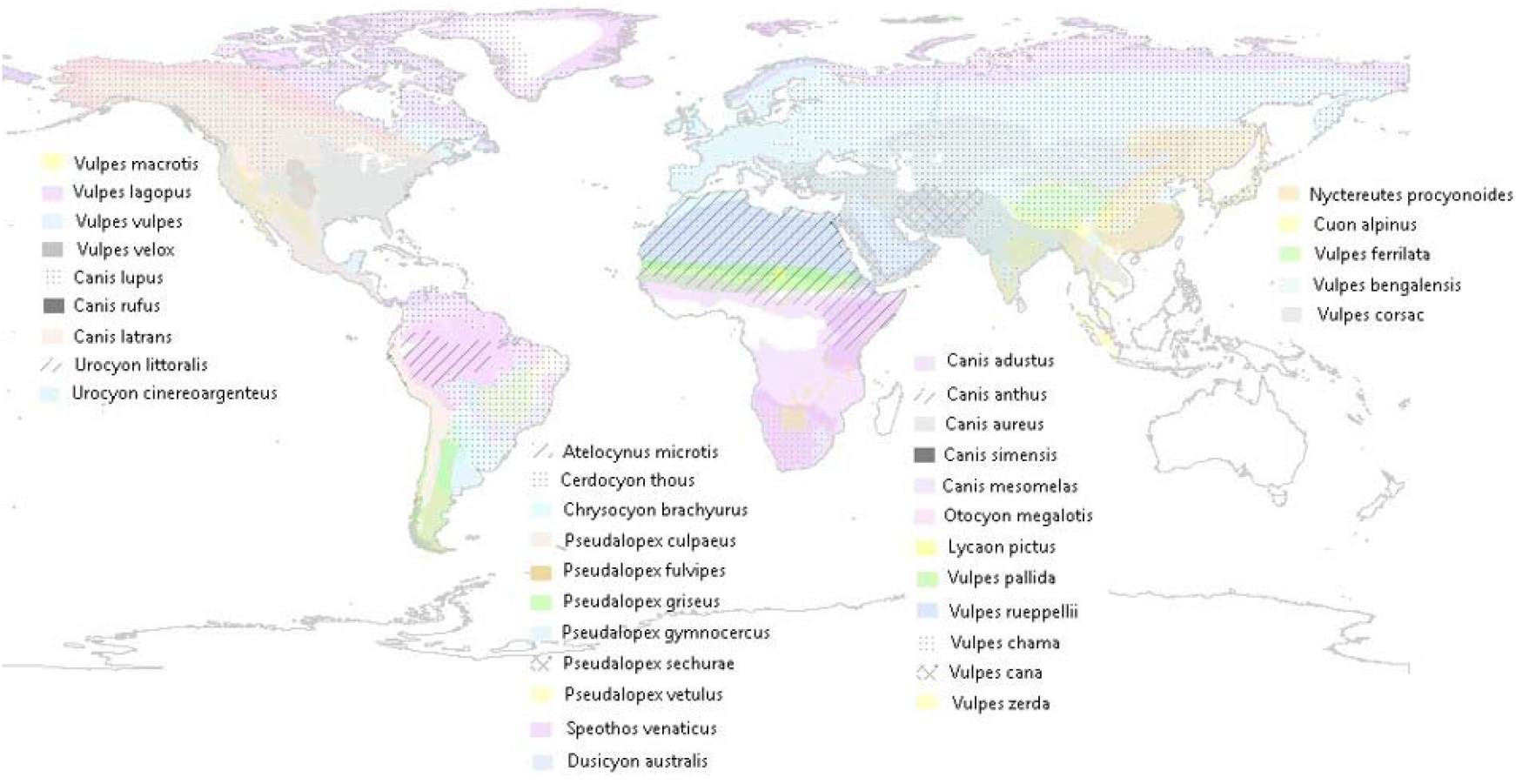
Distribution polygons used here for all the 37 species of canids obtained from IUCN (2020).

**Figure S3.**
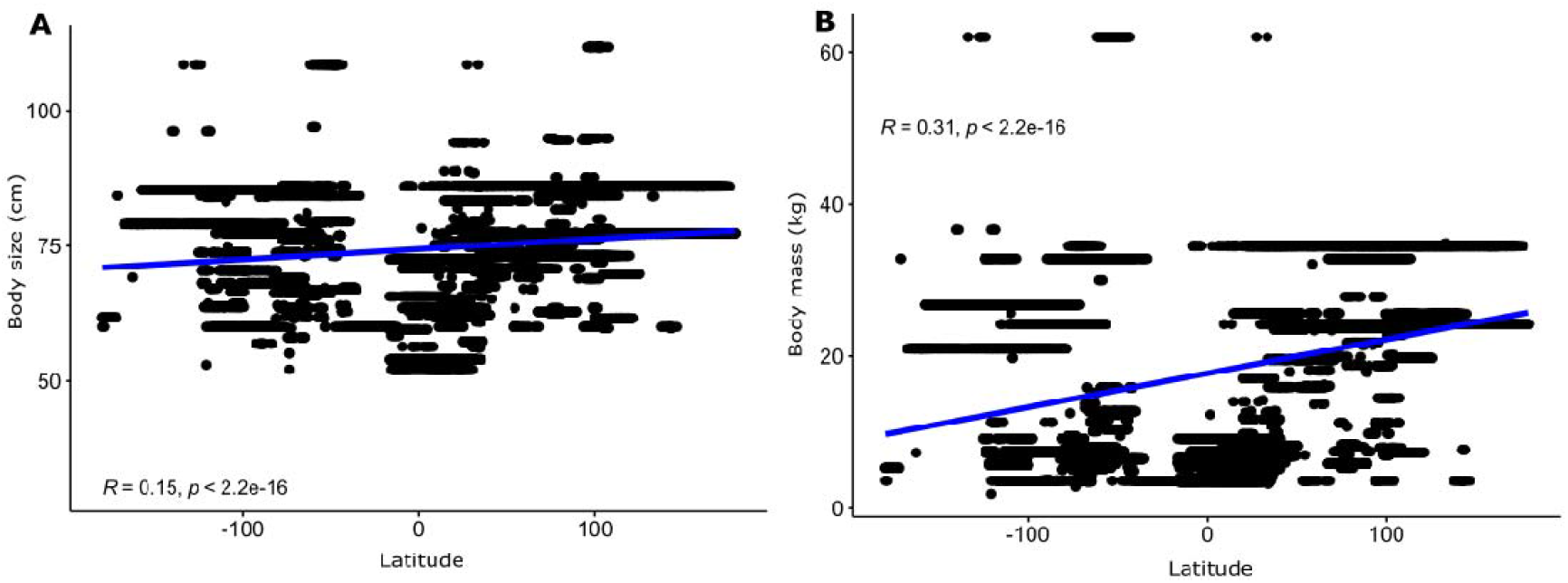
Relationship between the latitudinal gradient and traits that represent species’ sizes (A -body size and B -body mass).

